# Metagenomic surveillance of undiagnosed febrile illness in Nigeria does not reveal the etiological agent for most patients

**DOI:** 10.64898/2026.02.07.704564

**Authors:** Grace J. Vaziri, Julia C. Pritchard, Jillian I. Howard, Grace E. Stamm, David H. O’Connor, Christina M. Newman, Matthew T. Aliota, Asabe Dzikwi-Emennaa

**Affiliations:** Department of Veterinary and Biomedical Sciences, University of Minnesota, Twin Cities; Department of Pathology and Laboratory Medicine, University of Wisconsin-Madison.; Department of Veterinary Public Health and Preventive Medicine, University of Jos, Plateau State

**Keywords:** metagenomics, undifferentiated febrile illness, virus, etiology

## Abstract

Molecular and microscopy-based diagnostic capacity is often insufficient or unavailable in places where infectious disease burdens are highest, such as in West Africa. Rapid diagnostic testing (RDT) can provide quick and affordable diagnoses of common infections but is an imperfect solution due to limitations around detecting and dealing with false negative and false positive results. An alternative to RDT is unbiased metagenomic sequencing for pathogen surveillance. Here, we present data from unbiased metagenomic sequencing used to identify causes of undiagnosed febrile illness in Jos, Plateau State, Nigeria. Proof of concept for this approach has been demonstrated by several groups who have identified epidemic and endemic viral diseases like Lassa fever, yellow fever, and Chikungunya. Here, we show that unbiased deep sequencing and metagenomic analysis can be used to identify RNA viruses in clinical samples. We sequenced RNA from sera of patients (n = 343), many of whom were acutely febrile (76 %) in a survey of clinics in Jos. We detected five human-infecting viruses in 39 (11 %) specimens. Among these were hepatitis B virus, human pegivirus, and several anelloviruses. While most of the viruses identified are unlikely to cause clinical symptoms in the patients we sampled, their presence demonstrates the validity of our approach. Additionally, our sequencing data allowed us to identify genetic material from potentially pathogenic bacteria, another possible etiological agent of febrile illness.

## Introduction

In low– and middle-income countries, febrile illnesses with unknown etiologies are common while investigations of potential viral etiologies are rare^1,2^. For example, a review of studies investigating the sources of febrile illnesses noted that only 17 % of investigations even considered viral etiologies^2^. In Sub-Saharan Africa molecular diagnostics are limited, so rapid diagnostic testing (RDT) is often used to detect common fever-causing infectious agents, including malaria, typhoid, SARS-CoV-2, tuberculosis, and HIV^3–5^. The suite of potential causes of febrile illness, however, far exceeds the capacity for RDT, and includes fungal, bacterial, protozoal, and viral infections^5^. While RDT has been generally considered a success story in Africa, there are limitations, including sensitivity and specificity issues, cross-reactivity leading to misdiagnosis, and the inability of these technologies to characterize the infecting agent if a patient tests negative. Failure to detect and identify the true etiology of febrile illness has detrimental effects for both individual patients and public health^6^. Individual patients without a definitive diagnosis are at greater risk of receiving incorrect (or no) therapeutic interventions. Further, if clinicians are unable to detect the source of febrile illness, they may resort to the presumptive use of antimalarials and antibiotics in a stopgap effort to treat their patients. Inappropriate use of antimicrobial drugs is termed ‘overtreatment’ and can promote the evolution of drug resistance among their intended targets (e.g., the spread of mutations conferring artemisinin resistance in *Plasmodium falciparum* in Cambodia and Rwanda;^7–9^). A study of children under five years old in Nigeria documented an overtreatment rate of 83 % based on microscopic confirmation of malaria parasitemia^10^. Ultimately, failure to detect and identify sources of febrile illness may stymie the ability of public health workers to implement effective public health interventions and could potentially enhance the progression of infectious disease outbreaks to epidemic levels.

Metagenomic sequencing (mNGS) is increasingly used as a method for detecting the presence of pathogens from both individual-level (i.e., patient^11–13^) and aggregate-level (e.g., air, wastewater^14,15^) samples. The taxon-agnostic approach of metagenomic sequencing, in which all genetic material (RNA or DNA, depending on study goals) from a sample is sequenced and identified, distinguishes it from targeted approaches, in which samples are assessed for the presence of predetermined pathogens of interest. When causative agents of illness are unknown and potentially heterogeneous across a patient cohort, mNGS can be a powerful tool for identifying the presence of pathogenic microbes. The potential information gained from sequencing approaches is especially enticing in cases where RDT and patient symptomology yield negative and/or non-specific conclusions. For example, a mNGS study of sera from acutely febrile children in Tanzania identified rotaviruses and enteroviruses (both in family *Picornaviridae*) in >10 % of samples; these infectious agents aren’t traditionally monitored for using RDTs^16^. Beyond improving the ability of clinicians and researchers to identify potential fever etiologies, mNGS can detect the presence and circulation of asymptomatic infections. Identifying both asymptomatic infections and infections with relatively nonspecific symptoms is important to minimize cryptic transmission and exposure to susceptible patients for whom infections are likely to be more severe. For example, many arboviral pathogens, such as dengue, Zika, and West Nile viruses can sometimes result in asymptomatic infections, but in some patients, they can also cause severe pathology^17–19^. Early detection of these infections in both asymptomatic and symptomatic patients can improve the efficacy of public health responses.

In this study, we used mNGS to detect RNA from human-infecting pathogens among a group of patients who visited four clinics in Jos, the capital city of Plateau State, Nigeria in June-October 2023. Jos, Nigeria has a high burden of infectious diseases. Malaria and typhoid fever are common infections among patients from this region and often assumed to be the cause(s) of febrile illnesses^20^. We therefore collected serum from a mixed-sex, mixed-age population of patients that included febrile and afebrile patients. Samples came from patients with confirmed malaria or typhoid fever and from patients with no diagnosis. We report the detection of several common, nonpathogenic anelloviruses, as well as two detections of hepatitis B virus, and reads from 53 bacterial taxa known to infect humans.

## Methods

### Sample collection

Samples were collected from four clinics (Figure 1) during the 2023 rainy season (between late June and late October) in Jos, Plateau State, Nigeria. All samples were collected with permission from the Plateau State Ministry of Health (Ref. MOH/MIS/202/VOL.T/X issued 9th May, 2023). We obtained additional permission from two of the clinics, Faith Alive Foundation (FAFEC/08/38/0039 issued 24th May, 2023), and Plateau State Specialist Hospital (NHREC/09/23/2010b issued 19th April 2023), and the remaining two clinics, Comprehensive Health Centre Dadin Kowa and Famoseg, required no additional approval beyond that granted by the Plateau State Ministry of Health. A subset of patients were administered RDTs for either malaria (*P. falciparum*) or typhoid (Table 1). Data were collected from anonymized patients who agreed to participate in the study at the time of their clinic visit. Socioeconomic, demographic, and symptom-related data were collected by trained interviewers who obtained patient consent prior to reading questions aloud and recording patients’ responses in a standardized form. Serum was separated from blood collected during clinic visits and stored in 2 mL cryotubes at –20 °C for storage prior to export. Samples were picked up on 29 January 2024 from University of Jos and were transported from the Lagos airport to the Frankfurt airport where they remained until 3 February 2024 due to a strike among airport staff. Dry ice was reported to be replenished during this time, but the courier provided no confirmation of this. Samples were then shipped to New York, USA where they arrived on 4 February 2024 and remained until 19 February 2024. Dry ice was replenished daily during this period. Although shipping containers contained temperature monitors, all were nonfunctional when they arrived and requests for temperature reports from the courier were denied. Serum appeared to have remained frozen during transit (evidenced by tubes retaining serum in the bottom of the tube, despite being jostled and arriving upside-down), but we cannot definitively confirm that they remained frozen throughout all transit. Following receipt of samples in the United States, samples were stored at –80 °C until RNA extraction.

**Figure 1.**
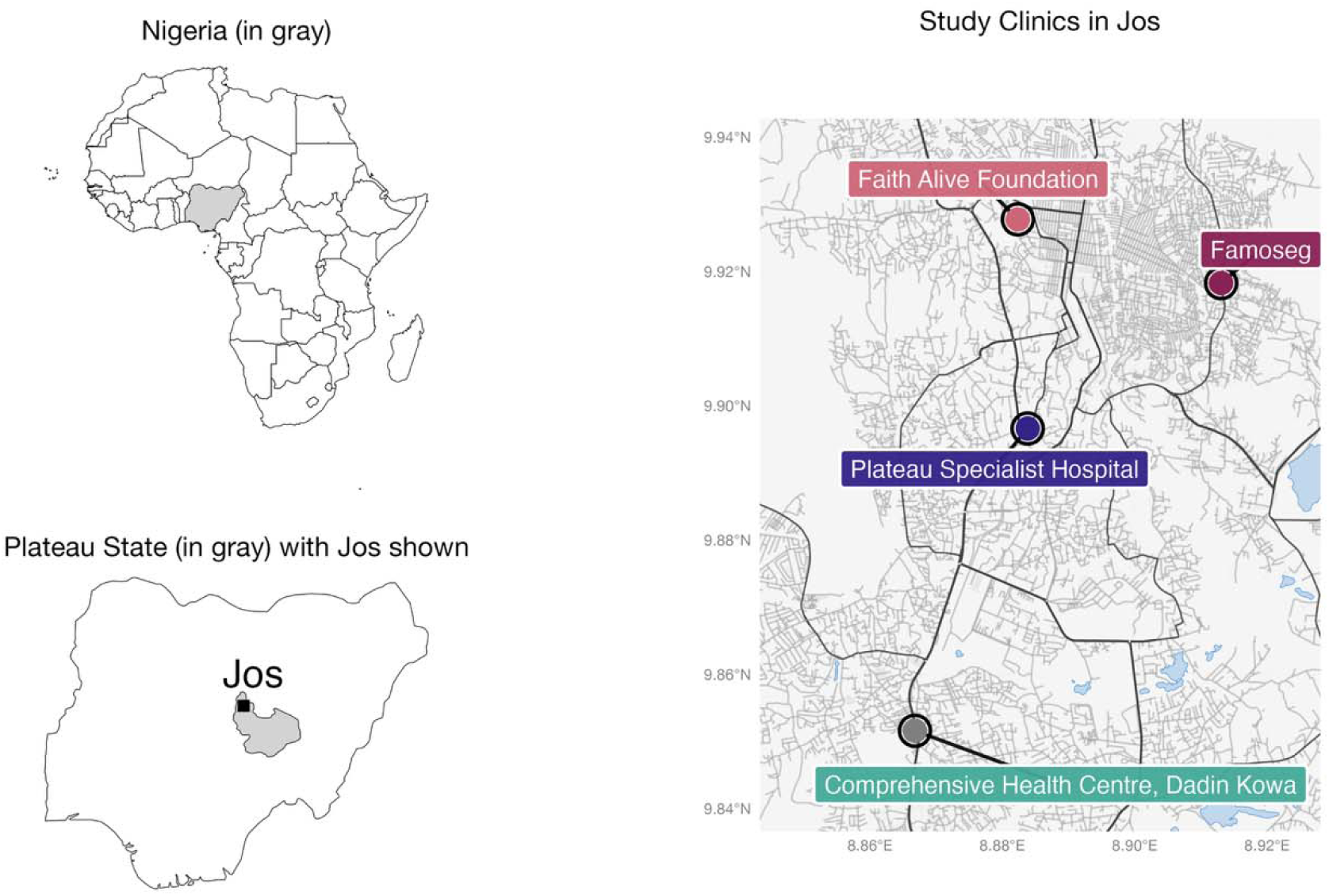
Serum samples were collected from patients who visited four clinics in the city of Jos, Plateau State, Nigeria. Sample collection was unevenly spread across hospitals, with most samples coming from the Faith Alive Foundation and Famoseg clinics. **Figure 1 alt text.** Three maps showing the continent of Africa with the country Nigeria in gray, Nigeria with Plateau State in gray and the city of Jos marked with a bullet point, and a street map zoomed in on Jos showing the locations of the four clinics from which samples were collected.

**Table 1.** A small subset of patients was confirmed to have (or to not have) malaria or typhoid infections, however, most were not provided with a diagnosis at the time of sample collection.

| Diagnosis | N (all) | N (NVD) | N (CZ ID) |
| --- | --- | --- | --- |
| Malaria negative, Typhoid positive | 15 | 8 | 7 |
| Malaria negative, Typhoid unknown | 152 | 119 | 111 |
| Malaria positive, Typhoid positive | 1 | 1 | 1 |
| Malaria positive, Typhoid unknown | 24 | 19 | 20 |
| Typhoid negative, Malaria unknown | 1 | 1 | 1 |
| Typhoid positive, Malaria unknown | 11 | 7 | 7 |
| No diagnosis | 282 | 188 | 212 |
| **Total** | **486** | **343** | **359** |

### Sample Processing

Prior to RNA extraction, samples were inactivated in a biosafety level three (BSL 3) laboratory with TRIzol LS supplemented with chloroform at 20 %, and then RNA was extracted at BSL 2 according to the manufacturer’s recommendations. RNA was re-suspended in a final volume of 30 µL nuclease-free water. A no template control sample (water) was extracted each time RNA was extracted from a batch of serum samples to control for contamination in the extraction space or reagents. To avoid confounding effects due to RNA extraction batch, sampling clinic, or time of sample collection, we ensured that clinics were proportionally represented in each extraction batch according to their prevalence in the total sample set, and then randomly assigned samples to RNA extraction batches. All RNA was stored at –80 °C immediately after extraction until cDNA synthesis.

### cDNA synthesis

Double stranded cDNA from extracted RNA transcripts was prepared using modified sequence-independent single-primer amplification (SISPA) approach^21–23^. To reverse transcribe RNA templates, we prepared a 40 pmol/μL working stock of primer A (5’-GTT TCC CAC TGG AGG ATA-(N9)-3’) and added 1 µL of primer A working stock to 4 µL of extracted RNA. We heated this at 65 °C for five min followed by five min of cooling at 4 °C. To perform first strand synthesis, we added 2 µL 5x RT Buffer, 1 µL 10 mM dNTPs, 1 µL nuclease-free water, 0.5 µL 0.1 M DTT, and 0.5 µL SuperScript IV RT enzyme to each sample (all reagents from Invitrogen 18091200) and incubated for 10 min at 42 °C. We performed second strand synthesis by adding 1 µL of Sequenase reaction buffer, 0.15 µL Sequenase DNA polymerase (Applied Biosystems 70775Y200UN), and 3.85 µL nuclease-free water to each reaction and incubating at 37 °C for eight min. After incubation, we added 0.45 µL Sequenase dilution buffer and 0.15 µL Sequenase DNA polymerase (Applied Biosystems 70775Y200UN) to each sample and incubated again for eight min at 37 °C. Finally, we amplified 5 µL of template cDNA by adding 2 µL of primer B (5′-GTT TCC CAC TGG AGG ATA-3′), 25 µL LongAmp Taq 2x master mix (NEB M0287S), and 18 µL nuclease-free water and running PCR with the following conditions: 94 °C for 30 s, followed by 30 cycles of 94 °C for 15 s, 47 °C for 20 s, and 65 °C for 2 min, followed by 65 °C for 10 min. We prepared a negative control during the cDNA synthesis step for each batch of cDNA synthesis using nuclease free water instead of an RNA template. Following cDNA synthesis, we quantified 2 µL of each cDNA product using the Qubit dsDNA HS Assay Kit (Invitrogen Q32851).

### Nanopore library preparation

We prepared libraries for nanopore sequencing using native barcoding (Oxford Nanopore SQK-NBD114-24 and SQK-NBD114-96) (https://nanoporetech.com/document/ligation-sequencing-gdna-native-barcoding-v14-sqk-nbd114-24?device=MinION). Briefly, we followed the protocol provided by Oxford Nanopore to take forward up to 400 ng of cDNA (or 11 µL of cDNA template if cDNA concentration was insufficient) into DNA repair and end preparation. After end preparation, we ligated native barcodes, pooled barcoded samples, and performed adaptor ligation and cleanup. Following adaptor ligation, we cleaned up the library with AMPureXP beads and washed the beads twice with 125 µL short fragment buffer. In addition to patient samples and negative controls from previous steps (RNA extraction and cDNA synthesis), each of our final sequencing libraries contained a negative control (nuclease-free water) on which we performed preparation and barcoding to control for contamination in the library preparation reagents and spaces. Additionally, we extracted RNA, synthesized cDNA, and sequenced at least two positive controls with every library. The first of these positive controls was a Powassan virus (POWV) isolate used in the Aliota (MTA) laboratory and included to ensure that our sequencing approach could successfully handle viral RNA. The second of the positive controls were synthetic RNA spike-ins (SRSIs) containing 10,000 gene copies of a foreign control sequence^24^. We included a unique SRSI (sequences in Supplementary Data 1) with each library we sequenced and used these sequences to track cross-contamination rates between sequencing runs.

We eluted the final library in 15 µL of elution buffer and quantified average fragment length and concentration using an Agilent Bioanalyzer. Using the average fragment length and concentration of our final library, we calculated the volume of library that comprised 300 fmol to carry forward to sequencing. Previous experience using nanopore sequencing for mNGS applications indicated that recommended loading concentrations from the product manual are too low for libraries, as they comprise generally short fragments (about 490 bp) leading to underloading of flow cells and reduced data yields.

### Bioinformatic methods

We basecalled all libraries using the high-accuracy basecalling model (dna_r10.4.1_e8.2_400bps_hac@v5.0.0). After demultiplexing libraries and discarding those with fewer than 10,000 raw sequencing reads, we processed samples using the *NVD* (v. 2.0) workflow (https://github.com/dhoconno/nvd). The *NVD* workflow performs initial classification of human virus sequences with the NCBI STAT tool^25^. Following STAT classification, the workflow uses a two-step BLAST verification, first performing a rapid sequence similarity search with megablast, then using the more sensitive blastn tool to align and classify contigs that were unclassified with megablast. The result of this two-step classification procedure was a list of all megablast hits for any contig that BLASTed to a viral reference sequence, and up to five blastn hits for any contig that blasted to a viral sequence. To generate a one-to-one list of contigs and taxon assignments (BLAST hits), we selected the single best match based on E score for each sample/contig combination. We imported this list into R (v. 4.2.2) along with a list of read counts per contig to generate a taxon-by-sample matrix filled with read-counts (i.e., an OTU table) with *phyloseq* v. 1.44.0^26^. Next, we used the ‘isContaminant()’ function, a statistical decontamination algorithm, from the *decontam* (v. 1.18.0) package^27^ with the ‘combination’ method enabled and specifying the sequencing library as a batch variable. After removing reads identified by *decontam* as statistical contaminants we removed reads from all non-viral taxa that remained in the dataset. We also consolidated reads from viruses that were uninformatively listed as being separate (e.g., reads from Human pegivirus and *Pegivirus hominis* were combined). While *NVD* is specifically designed to detect viruses that can infect humans, the method of taxonomic classification (megablast + blastn) combined with subsequent selection of a single best taxonomic assignment led to non-viral reads being included in the dataset. This occurred when a contig did not unanimously blast to viral sequences and a non-viral sequence had the lowest E-value of all blast hits. Finally, we manually inspected the remaining taxa to verify they met the following criteria: 1) Are known to infect humans, 2) Are not possible lab contaminants, 3) Have at least 5 reads.

In addition to our virus-focused analysis, we used two methods of taxon-agnostic sequence profiling: *gottcha2* (v. 2-2.1.10) and CZ ID (formerly IDseq). The *gottcha2* pipeline is a gene-independent taxonomic profiling algorithm incorporated into the larger *NVD* workflow. *Gottcha2* classifies bacterial and eukaryotic reads, in addition to viral^28^. We first filtered to only retain samples with at least 10,000 raw sequencing reads. Then, we specified the following *gottcha2* parameters to accommodate our relatively low biomass samples: cutoff percent of 0.0001(allowed us to detect very rare taxa), taxa stringency of 0.5 (allowed for relatively low confidence requirements), entropy of 0.7 (allowed for more repetitive sequences), per-sample contigs contained at least 100 consecutive bases, and at least 100 reads. Finally, we disabled quality trimming, which is less critical for Nanopore data. We processed the output of the *gottcha2* profiling similarly to how *NVD* output was processed with several modifications. First, as previously described for *NVD* –generated data, we performed statistical decontamination using the ‘combined’ method of the ‘isContaminant()’ function in package *decontam*. Next, we used the ‘ConQuR’ function (with penalized settings) from package *ConQuR* (v. 2.0) to detect and mitigate batch effects in our data^28^. The ConQuR algorithm accounts for within-batch variability in library size and assumes that batch and mean library size are confounded, allowing the use of raw (non-normalized or rarefied) libraries as input data^29^. Our samples went through three ‘batched’ processes: RNA extraction, SISPA, and sequencing library preparation. To identify which of these processes imposed the greatest batch effect, we first conducted a permutational analysis of variance (PERMANOVA) using the ‘adonis2’ function in package *vegan*. After determining that RNA extraction batch accounted for the largest amount of variation (R2 = 9.9%, p < 0.005), we proceeded with batch correction based on RNA extraction batch. To select an appropriate “reference” batch against which to correct the batch effects of other RNA extraction batches, we tested each RNA extraction batch as a reference batch and calculated the reduction in proportion of variation attributable to batch and the preservation of variation due to covariates (diagnosis, age, fever, cumulative number of symptoms, and clinic visited) after batch correction (Table S1). We did not perform batch correction for NVD-generated virus data because the filtering steps resulted in a dataset too sparse to detect batch effects. Following batch correction, we subset the bacterial dataset according to the following criteria: 1) Retain reads from bacterial species included in a comprehensive list of bacterial pathogens infecting humans^30^, 2) Taxon must have at least 10 reads in a sample, and 3) Taxon must have at least two contigs in a sample.

To complement our *gottcha2* results, we also uploaded raw fastq files to the CZ ID (formerly IDseq) nanopore pipeline^31^. CZ ID is a cloud-based, open-source taxon agnostic bioinformatics pipeline that downsamples to a maximum of 1 million nanopore reads per sample. After running the pipeline, we generated a Z-score metric to distinguish reads due to contamination from potential pathogens or commensals in our samples^32^. Z-score was calculated as the difference between a taxon’s reads in a biological sample and the taxon’s mean number of reads in control samples, divided by the standard deviation of the taxon’s reads in the control samples. We filtered our dataset to include taxa with Z-scores >= 1 and alignment length (sample reads against reference genomes) of at least 200 nucleotides, and to include those represented by at least 10 bases per million in our sample set and at least two reads per sample. We also discarded alignments <= 90 % similarity to reference alignments, and those with alignment significance (E values) > 10-100 (the closer an E value is to zero, the more significant the alignment).

### Corroborating RDTs with mNGS

We investigated whether our mNGS approach could identify infections diagnosed with RDTs. For both malaria (caused by *Plasmodium* spp.) and typhoid (caused by *Salmonella enterica* serovar Typhi [*S.* Typhi]), we categorized samples based on their RDT diagnosis (positive, negative, unknown) or their sequencing results (whether, after applying filtering criteria described above, samples had reads classified as *P.* (*vivax, falciparum, knowlesi, malariae* or *ovale*) or *S.* Typhi).

To investigate discordance between RDT and sequencing results for malaria detection, we mapped all reads from samples across RDT/sequencing concordance categories against six *Plasmodium* reference genomes (three *P. falciparum* assemblies, and one assembly each for *P. knowlesi, P. vivax,* and *P. ovale*) using minimap2 (v. 2.30). To validate whether observed alignments represented true *Plasmodium* infections, we selected representative samples from each RDT/sequencing concordance category (n = 6 samples) and the sample with the highest alignment rate against any *Plasmodium* genome (n = 1 samples) and performed taxonomic classification using Kraken2 (v2.1.2) with a protozoa-based database. Reads classified as *Plasmodium* by Kraken2 were then verified using BLASTn against NCBI’s nt database to determine their true taxonomic origin. We repeated the second half of this workflow for all samples that were RDT– or sequence-positive for *S.* Typhi. Briefly, we performed taxonomic classification of sequences from these samples with a Kraken2 database (bacteria build) and then pulled and blasted all reads classified by Kraken2 as *S.* Typhi. We bypassed the minimap2 step used in our *Plasmodium* spp.-focused investigation because the high serovar diversity within the subspecies *Salmonella enterica enterica* would make mapping reads to a single genome within the subspecies complex uninformative (reads from non-Typhi serovars are also expected to align at high rates to the *S.* Typhi genome).

### Data availability

The sequencing data generated in this study have been deposited in the Sequence Read Archive (SRA) under bioproject ID PRJNA1417234.

## Results

### Patient population

Samples were collected at the clinics from 486 patients. The population represented in this subset of the samples ranged in age from 2 to 75 (median age = 28) and comprised a sex ratio of approximately 1:2 M:F (155 male, 294 female, and 32 patients for whom sex was not recorded). Of these patients, 78 % (377/486) presented to clinics with fever, and 74 % (358/486) had, in addition to fever, at least one of the following symptoms: hematuria (bloody urine), difficulty urinating, rash, jaundice, diarrhea, bloody diarrhea, vomiting, joint pain, headache, myalgia (muscle aches), conjunctivitis, malaise, skin hemorrhage, bleeding gums, nosebleed, or edema. At the time of sample collection, 25 patients included in the study tested positive for malaria and one malaria-positive patient also tested positive for typhoid, the disease caused by the bacteria *S.* Typhi. Twenty-seven patients (including the coinfection case) tested positive for typhoid (one negative), and 282 patients received no diagnosis (or no diagnosis was recorded, Table 1). Patients were not evenly spread across the four clinics included in the study, rather 15 % (72/486), 47 % (226/486), 32 % (156/486), and 7 % (32/486) came from Comprehensive Health Centre Dadin Kowa, Faith Alive Foundation, Famoseg, and Plateau Specialist Hospital clinics, respectively.

### Sequencing performance

We prepared sequencing libraries and sequenced 417 patient samples, as well as 20 cross contamination controls, 56 no template controls, and 19 positive control samples (n= 512 sequencing libraries total). Different subsets of sequencing libraries generated from patient samples were retained for different analyses, depending on analysis-specific filtering criteria. We achieved variable library sizes due to different levels of multiplexing, sequencing runtime, and sample quality (amount of host contamination varied from sample to sample). After discarding libraries with fewer than 10,000 raw reads, we retained 343 sequencing libraries (range = 108,999 to 5.2 M, mean (SD) = 1.2M (801,293) reads/library) for *NVD* analysis. The samples comprising these libraries were representative of the full patient dataset (median age = 28, approximate sex ratio 1:2 M:F, 76% febrile, 72% having fever and one additional symptom). Libraries with greater sequencing depth were more likely to contain reads assigned to human viruses (*R*^2^ = 0.97; Figure 2). Reads assigned to human viruses were relatively rare, however, comprising 9.0 % of all reads in samples (Figure 3). Cross contamination rates among sequencing batches were low (mean = 0.15 %, median = 0.0005 %).

**Figure 2.**
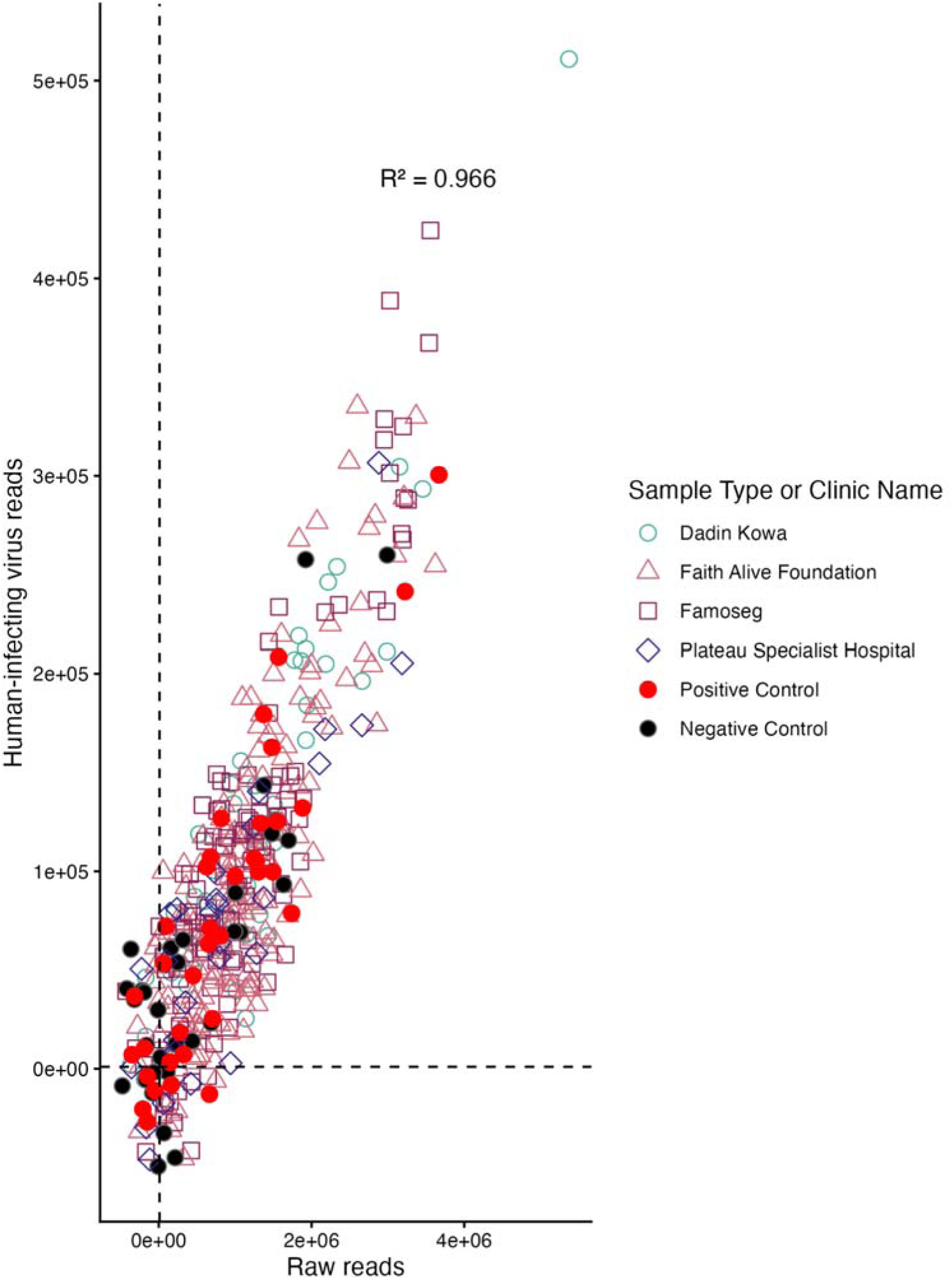
Sequencing depth was strongly associated (*R*^2^ = 0.97) with the ability to detect reads from human-infecting viruses. Samples to the right of and above the dashed vertical and horizontal lines contained at least 10,000, and 1,000 raw reads and reads from human-infecting viruses, respectively. **Figure 2 alt text.** Scatterplot showing the correlation between the number of raw sequencing reads on the x axis and the number of reads from human infecting viruses on the y axis. Solid points shown with circles are control samples (black = negative, red = positive), and open shapes represent the four clinics included in the study. Dashed lines indicate samples with more than 10,000 raw reads (vertical) and 1,000 reads from human-infecting viruses (horizontal).

**Figure 3.**
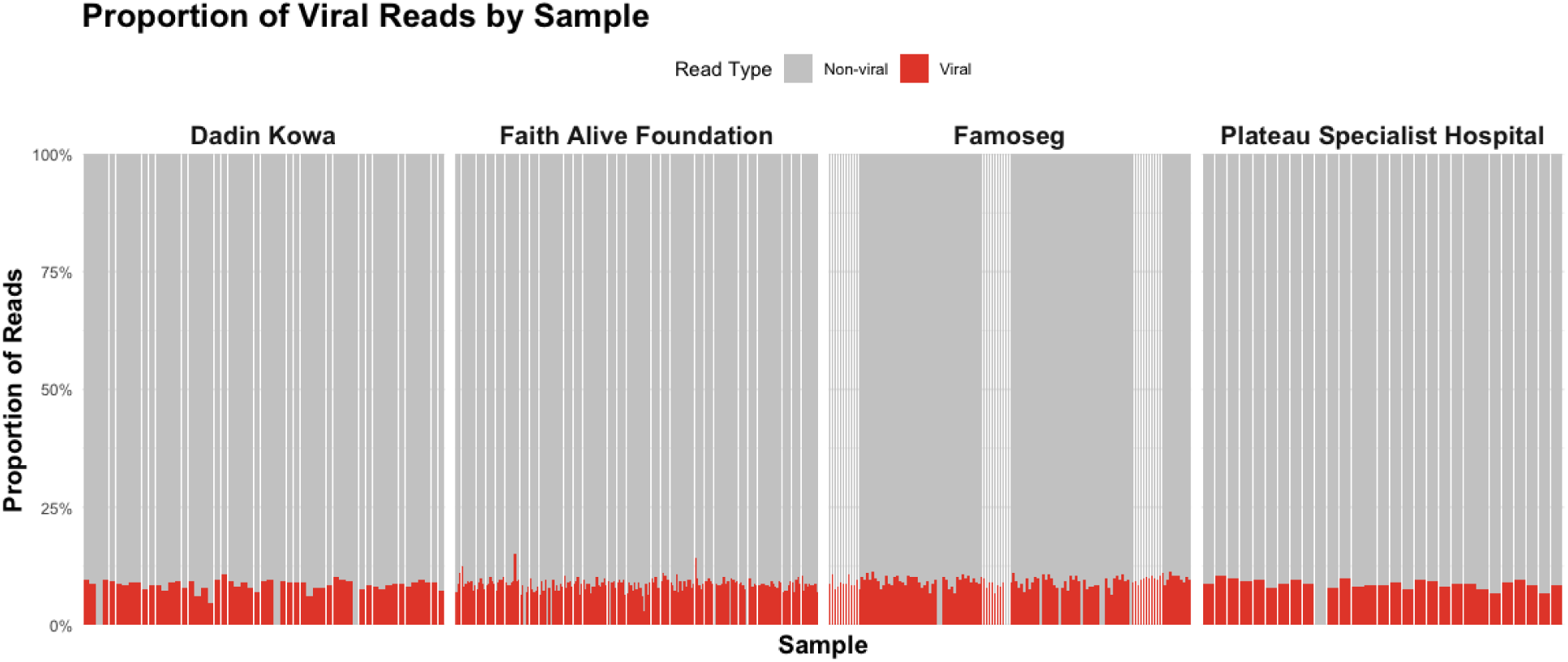
Prior to statistical decontamination NVD assigned a small proportion (mean = 8.4%) of raw sequencing reads to viruses known to infect humans. **Figure 3 alt text.** Stacked bar charts showing proportions of reads from viral (red) versus non-viral (gray) sources. Each bar is a sample, and samples from different clinics are separated on the x axis.

### Viruses

After filtering reads according to the criteria described previously, we retained six viral taxa in 39/343 samples (Figure 4), representing only 11 % of the patient samples with usable sequencing data. We identified one known vertebrate virus, hepatitis B virus (HBV, n = 2), as well as five human viruses not currently believed to cause disease: torque teno virus (TTV, n = 11), TTV-like mini virus (n = 19), human pegivirus (*Pegivirus hominis,* HPgV, n = 8), hepatitis delta virus (n = 2), and species in the Gammotorquevirus genus (n = 3). The most frequently detected virus was TTV-like mini virus in 5.5 % (19/343) of samples that produced usable sequencing data. The virus with the highest read count in one sample was HPgV in sample FA5, where there were 72,425 reads classified as being from HPgV. The HPgV reads recovered from this patient’s sample covered approximately 76 % (average depth 37 x) of an HPgV genome isolate identified in Nigeria in 2018 (Figure 5A). Prevalence of HPgV in our sample set was low (∼2.3 %, 8/343). We detected sequences from HBV in samples from two patients (Figure 4). In one patient, DK13, the 3,235 reads classified as HBV aligned with > 98 % similarity (and with equal coverage) to HBV isolates from other patients from Nigeria and Cameroon^33–35^ (Figure 5B). In the second patient with HBV reads, FMS115, the 5,552 reads mapped with > 98 % similarity to an HBV sub-genotype A clone generated by a South African research group (Genbank accession PP790594.1; Figure 5C). Both patients with reads classified as HBV were febrile and reported having headaches, and patient DK13 additionally reported joint pain and myalgia. Data on malaria and typhoid positivity status were unavailable for both patients.

**Figure 4.**
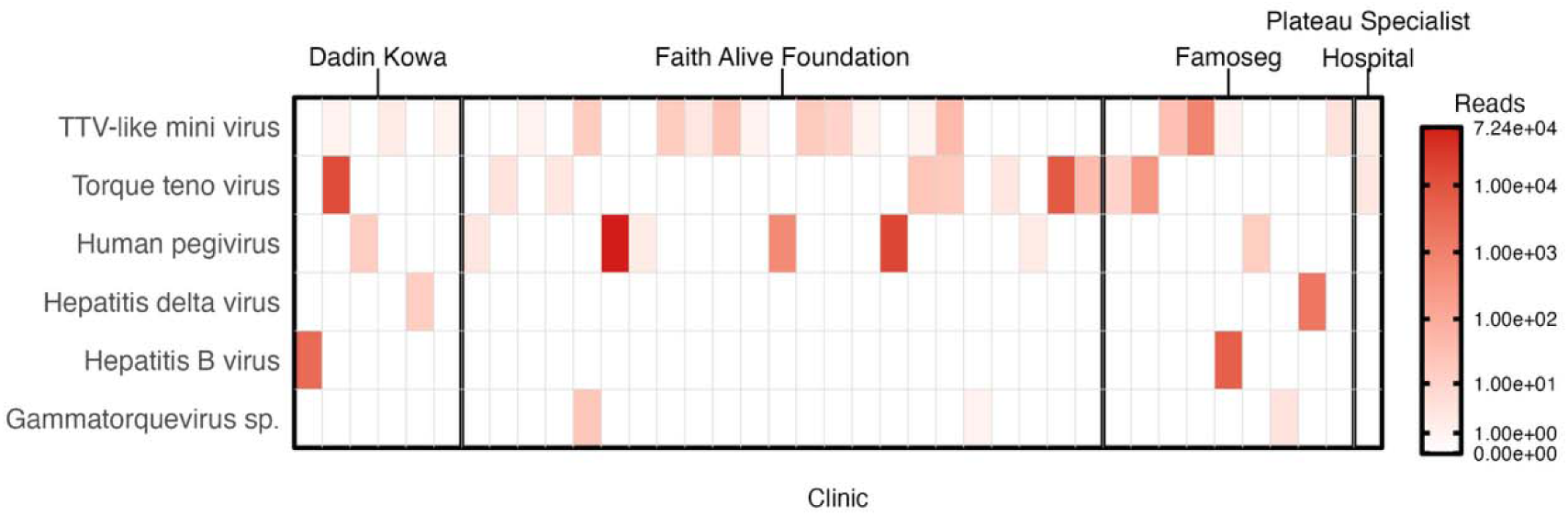
Most human-infecting viruses detected were non-pathogenic anelloviruses. Two patients had reads from hepatitis B virus. Columns representing individual patients are grouped by clinic. Box color saturation indicates the number of reads classified as each virus (rows) with more saturated colors corresponding to more sequencing reads for a virus x patient combination. Only patients with human-infecting virus reads are shown. **Figure 4 alt text.** Heatmap of viral read abundance in patient serum samples separated by clinic.

**Figure 5.**
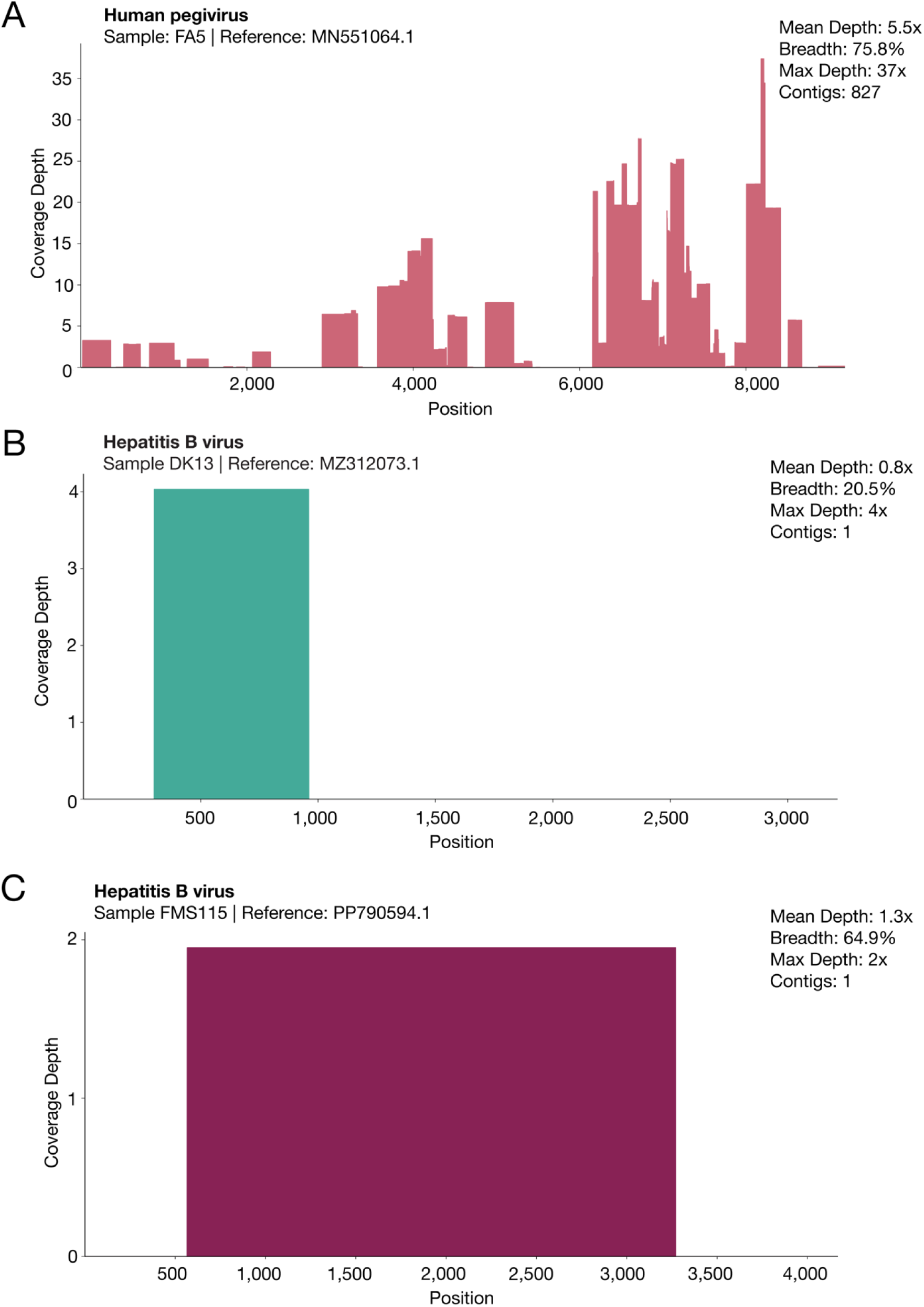
Representative coverage plots illustrating genomic coverage of (A) Human pegivirus reads recovered from sample FA5, (B) Hepatitis B virus reads recovered from sample DK13, and (C) Hepatitis B virus reads recovered from sample FMS115. Reference genome accessions against which reads were mapped are listed in plot subtitles. **Figure 5 alt text.** Three bar charts with bar height corresponding to number of reads recovered from a sample, and x axis position corresponding to the genomic location of reads on the most-closely related reference genome from BLAST. The top bar chart has a y axis that extends to 40 reads, the middle bar chart has a y axis that extends to 4 reads, and the bottom bar chart has a y axis that extends to 2 reads. Bar chart color corresponds to clinic identity.

### Taxon-agnostic analysis results

We used different filtering criteria to parse our *gottcha2*-derived results and our CZ ID-derived results, following general recommendations from other groups who have performed mNGS studies for our CZ ID filtering. Prior to applying the ‘potential pathogen’ filter, our CZ ID dataset contained 359 samples (Table 1). After filtering to retain only ‘potential pathogen’-flagged taxa, our dataset contained 315 samples and 83 taxa, of which 4 were viral, 73 were bacterial, and 6 were eukaryotic. Two samples (FMS159 and F17) that were successfully analyzed by NVD repeatedly failed to pass through the CZID analysis pipeline and were discarded. For parsing *gottcha2* data, we used filtering and decontamination criteria described in the methods, retaining only bacterial taxa with the potential to infect humans. This resulted in a dataset of 52 samples and 53 bacterial taxa (Table S2), which we further investigated for potential bacterial infections.

Bacterial taxa remaining in this dataset were represented in each sample by at least two contigs and 10 reads and were classified as species included in a published list of bacterial pathogens infectious to humans^30^. Read depth and breadth of human-infectious bacteria were relatively low; the maximum number of reads assigned to any single taxon in a sample was 20 and median coverage breadth (positions of a reference genome recovered in our sequencing data) was 2,500 bases. The most frequently detected bacterial species in this reduced dataset was *Enterobacter kobei*, which we observed in 12/343 (3.5 %) samples. This taxon also had the greatest coverage in our dataset; in one sample (FMS116), *Enterobacter kobei* reads covered 14,665 positions (0.3 % of the 4.8 Mb genome) of the two reference genomes to which our reads BLASTed.

### Plasmodium and Typhoid RDT concordance

Decontamination filtering of our *gottcha2*-derived data classified all *Plasmodium* spp. reads as contamination; all references to *Plasmodium* parasites detected with sequencing data going forward pertain to alignments that passed CZ ID filtering steps (described in methods). Among samples from patients with recorded RDT diagnoses for malaria (n = 21 positive, 119 negative), we found poor concordance between RDT and sequencing results (Table S3). Of the 21 patients who tested malaria-positive via RDT, only one (FA10) also had reads that passed our filters and mapped to *P. falciparum* in CZ ID. Among the 119 patients who tested negative for malaria via RDT, we detected filter-passing reads from *P. falciparum* in five samples.

Alignment rates to *P. falciparum* (3D7) references were uniformly low across all sample categories, ranging from 0.01 % to 1.22 % with no clear distinction between RDT-positive and RDT-negative samples. Samples showed substantially higher alignment rates to *P. knowlesi* (0.1 % to 42.8 %) and *P. vivax* (0.09 % to 19.4 %) references compared to *P. falciparum*, with this pattern persisting even in RDT-negative control samples.

BLAST validation of Kraken2-classified *Plasmodium* reads from representative samples (one per RDT/sequencing concordance category) showed that these sequences were predominantly human (84 %) or other primate sequences (9-16 % *Macaca*), with zero reads confirmed as authentic *Plasmodium* species. Investigation of FAL2, an RDT-negative sample showing the highest alignment rate to *P. knowlesi* (42.8 %), revealed that reads mapping to the apicoplast genome were bacterial sequences (*Achromobacter* spp.), confirming that high alignment rates resulted from contamination rather than true *Plasmodium* infection. This pattern was consistent across all tested samples regardless of RDT or sequencing status, indicating that the observed minimap2 alignments to *Plasmodium* genomes represent non-specific mapping of human, primate, and bacterial reads rather than true malaria parasite signal.

## Discussion

We used metagenomic sequencing of RNA from human serum samples to detect potential causes of febrile illness. Definitive diagnosis of the infectious etiology of a patient’s symptoms is beyond the confirmatory abilities of the data generated with mNGS techniques. Under ideal circumstances, however, such data could be used to narrow the range of or even pinpoint an etiological culprit which might then be confirmed with culture-, antigen-, microscopy-based, or targeted molecular diagnostic testing. The data included in the present study represent a culmination of several suboptimal, but not uncommon, circumstances that may have influenced our ability to detect potential microbial sources of febrile illness. These circumstances, including uncertainty and complications surrounding sample storage conditions during export, as well as host and laboratory contamination, may have constrained our ability to isolate, sequence, and identify RNA from every potentially pathogenic microbe in the dataset. Substantial variation in sequencing depth has been reported in previous mNGS studies of clinical samples, ranging from thresholds as low as 1,000 reads/sample to at least 2 million reads/sample. Our average sequencing depth of 1.2 million reads per sample falls within this range and is consistent with other recent studies in which negative results were common^12,13^.

Sample quality in our study may have been compromised prior to RNA extraction. During export and transport from Nigeria to the United States, the courier tasked with shipping samples experienced delays, during which time the cold chain of sample storage may have been disrupted. This suspicion is impossible to determine unequivocally, however, the sparse and relatively fragmented microbial nucleic acid content extracted from our samples suggests that some sample degradation may have occurred during transit. Independent of issues related to sample transit and storage, our samples overwhelmingly yielded host (human) reads. This issue is common among metagenomic pathogen surveys from human tissues^36,37^, especially from low biomass sample materials such as serum. The percent of reads in our study classified as being from viruses potentially pathogenic to humans (8.4 %) fell within the range reported by Grundy *et al.* (2023). While our analysis using *NVD* essentially performs *in silico* host depletion by white-listing for reads that come from human-infecting virus families, we did not perform any host-depletion steps during wet lab sample preparation. Other investigators have successfully dealt with host read contamination by performing a host-depletion process on patient samples^38^, however, including such a step is not without tradeoffs. Every additional procedure to which samples are subjected increases the potential for ‘kit-ome’ contamination, and imposes additional requirements in labor, time, expense, and expertise, which may make projects less feasible in resource-limited settings^39^. Metagenomic sequencing libraries of nucleic acid from low biomass samples, such as serum, are extremely susceptible to non-host contamination as well^39^. To counter this, we used negative controls at every step of laboratory work to control for contamination of reagents, and workspaces. Unfortunately, we could not control for potential contamination at the clinic level. Our *gottcha2* analysis PERMANOVA estimate indicated that there was a strong effect of clinic on the bacterial reads detected in patient sera. It is possible that patients from different parts of Jos, who presumably visited the clinics closest to their dwellings, do have relatively distinct microbial communities, but it is much more likely that the clinics themselves varied in the suite of contaminants to which samples collected at those clinics were exposed.

Despite methodological challenges, we did detect several viral taxa that, based on other mNGS surveys of human sera, we were unsurprised to find in our samples. For example, others have frequently detected pegiviruses in metagenomic surveys of human serum^11–13,36^. We detected HPgV in 8/343 (∼2.3 %) samples with usable sequence data. HPgV is a common, apparently nonpathogenic virus that can cause long lasting high-titer viremia but links to disease are unsubstantiated^40^. While we do not interpret detections of HPgV to be informative from the standpoint of identifying an etiological source of disease, we do consider its detection to be an indication that our methodological approaches were effective, when sample quality was adequate and nucleic acid from pathogens was present in sufficient quantities to detect.

Our conclusions regarding the possible presence of bacterial pathogens in our samples are less certain. Even after employing the same laboratory control scheme, and even more stringent *in silico* filtering and decontamination methods as were used for data on viral reads, the bacterial reads we most often detected in our samples were infrequent human pathogens. The most commonly detected bacteria, *Enterobacter kobei* was seen in 12 samples (3.4 % of samples with usable sequencing data). In patient FMS116, *E. kobei* was represented by two contigs that BLASTed to two different reference genomes. Neither of these genomes were Nigerian in origin, and only one was isolated from a human infection (the other was isolated from an environmental source). Additionally, most published reports of *E. kobei* describe nosocomial infections, but our study did not collect the detailed health histories needed to assess this likelihood^41,42^. Our bacterial reads are difficult to interpret definitively, as uncommon bacterial infections are rarely diagnosed, and even more rarely sequenced and archived. While bacterial infections, especially *Salmonella*, are a leading cause of febrile illness in Sub-Saharan Africa^43^, we emphasize the importance of interpreting detections of human-infecting bacteria cautiously. On the flip side, given the sample-related challenges described previously, it is difficult to know whether ‘negative’ sequencing data (samples in which no fever-causing pathogen RNA is detected) truly represent the absence of pathogens, or simply the failure to detect them.

We leveraged the heterogeneity within our patient data with respect to RDT diagnosis for malaria and typhoid by conducting a concordance analysis. Specifically, we investigated whether our sequencing approach detected sequences from human-infecting *Plasmodium* species, or from *Salmonella enterica enterica* serovar Typhi (*S.* Typhi), in samples from patients who tested positive for infections with these pathogens via RDT. In all cases, we concluded that our sequencing did not produce sequences from these pathogens, however, had we treated initial CZ ID or *gottcha2* results as final, bypassing additional blastn checks and stringent contamination filtering and batch effect mediation, respectively, we would have considered both these infections to be prevalent in our sample set. We stress that failure to detect sequences from these infectious agents in samples from patients with known positive infection status does not nullify the validity of RDT results. Rather, this outcome is indicative of biological characteristics of the parasites, and limitations of our study design. As intracellular parasites, both *Plasmodium* spp. and *S.* Typhi are unlikely to be present in the cell-free sample material (serum) used in our study.

Researchers have successfully used mNGS to identify etiological culprits across different patient populations, sample types, and hospital settings (inpatient versus outpatient). For example, using cerebrospinal fluid of pediatric patients with idiopathic meningitis in Bangladesh, Saha and colleagues (2019) identified previously undetected neuroinvasive Chikungunya virus infections^44^. In Edo State, Nigeria, a rash of patients with unexplained febrile illness who presented to a clinic in the Lassa-endemic region were found to be infected with yellow fever virus using mNGS to a comparable sequencing depth (1.6 M reads/sample) to that used in our study (1.2 M reads/sample)^45^. Unlike these examples, in which mNGS was deployed in response to unexplained public health phenomena, our study is more representative of a routine surveillance scenario where the data could then be used to identify which pathogens to prioritize for diagnostic capacity and development of molecular assays. Our study was conducted on a cross-section of the patient population visiting four clinics in the capital city of Plateau State, Nigeria in 2023. Rather than responding to a surge of unexplained febrile illnesses in a demographic or temporal focus, our samples were collected opportunistically as patients visited clinics of their own volition to seek care. Therefore, despite the geographical proximity from which they originated, the samples used in our study are heterogeneous in several potentially consequential dimensions. Age, sex, and prior infection status all varied in our patient cohort. Our samples came from ambulatory outpatients who were able to get themselves (or be taken by their caretakers) to clinics, introducing another potential source of sample heterogeneity: the elapsed time between symptom onset, clinic visitation, and sample collection. Without standardization of the timing of sample collection, fever-associated viremia or bacteremia could have cleared prior to sample collection. In a different study of febrile patients in southern Nigeria, mNGS only detected dengue virus (DENV) reads in ∼58% of serologically-positive DENV samples tested, indicating that even confirmed infections are often not detected with mNGS^46^.

Our findings lend insight to several important considerations for mNGS investigations and researchers. During our investigation we were several times reminded that the exquisite sensitivity of mNGS can be a double-edged blade. On one hand, we show that despite the microbial sparsity and low biomass of the sample type we used (serum) and the minimal processing (intended to make our approach easy-to-deploy in low-resource settings) we applied to our samples, we were able to detect the presence of viruses known to infect humans. On the other hand, we also detected insidious lab-based contamination at very low (below qPCR detectability) levels from pathogenic viruses with endemicity to our study area. Specifically, we detected RNA from West Nile virus (WNV) and DENV in several samples. These viruses are known to circulate in Nigeria and are also among the viruses studied by researchers in the same physical space where we conducted the lab work for this study. Even after statistical contaminant removal and nucleotide BLAST searches on the sequences classified as WNV and DENV, ambiguity remained surrounding their origins (was their presence due to lab contamination or were they truly present in patient samples?). To answer this question with greater confidence, we used a tiled PCR (PrimalSeq) approach, following the methods of Quick *et al.* (2017) and Vogels et al. (2024), to attempt to recover and sequence the entire viral genomes of WNV and DENV from putative-positive samples (See supplemental methods)^47,48^. This approach allowed us to definitively rule out the possibility that the sequences were true positives (See supplemental results).

Despite diligent protocols for cleaning shared lab spaces, the use of negative controls at every step of the laboratory work, and employing post-sequencing bioinformatic methods for contamination removal, we found that trace amounts of fragmented RNA can still contaminate mNGS pipelines, potentially leading to false positives. Additionally, the labor and time required to rule out such spurious positives is considerable, suggesting that ideal implementations of mNGS approaches will include dedicated facilities, which may often be unrealistic. Given these challenges, it is perhaps unsurprising that the sequencing we performed primarily detected the presence of nonpathogenic commensal viruses (after controlling for potential contaminants). In fact, we suggest that the sensitivity of mNGS to timing of sample collection, susceptibility to contamination at each stage of the process, and required expertise for method implementation may make it more appropriate as a surveillance tool for samples collected in aggregate settings such as air and wastewater, rather than individual patient samples.

## Conclusion

Our goal in this study was to use mNGS to detect and identify possible causative agents of febrile illness among patients who presented to four clinics in Jos, Plateau State, Nigeria. We were able to detect common human-infecting viruses unlikely to cause fever, but only in a small subset of the samples we considered. Potential bacterial sources of fever and other symptoms were more commonly detected; however, coverage of bacterial genomes was low and symptoms due to bacteremia would need to be confirmed with follow-up cultures or qPCR. We suggest that in aggregate, public health monitoring settings, long term surveillance data generated with mNGS could provide valuable baseline estimates of expected microbial communities, allowing researchers and public health workers to flag potential pathogens of concern. We also recognize the utility of mNGS technology as a method for narrowing the range of potential etiologies of disease or pathology outbreaks in well-defined patient populations. Without capacity to provide a secondary, definitive diagnosis, the time, expertise, and resources required to carry out mNGS investigations may make them unsuitable as one-off additions to the clinician’s diagnostic toolkit, especially in resource-limited settings or when other patient data are sparse.

## Supporting information

Supplementary Methods and Results

## Acknowledgments

The authors acknowledge the University of Minnesota Genomics Core CoLab (especially Aaron Becker), for providing nanopore library prep reagents, and access to instruments and training.

## Financial Support

This work was supported by the Fulbright African Research Scholar Program (ADE), and NIH award R01 AI183302 (MTA).

## Authors’ current addresses

^1^Department of Veterinary and Biomedical Sciences, University of Minnesota, Twin Cities.

^2^Department of Pathology and Laboratory Medicine, University of Wisconsin-Madison.

^3^Department of Veterinary Public Health and Preventive Medicine, University of Jos, Plateau State.

## Notes

### Competing Interest Statement

The authors have declared no competing interest.

