## Supplementary Methods and Results for "Metagenomic surveillance of undiagnosed febrile illness in Nigeria does not reveal the etiological agent for most patients"

Supplemental methods

*PrimalSeq and DengueSeq*

For samples in which we detected reads aligning to the West Nile virus (WNV) or dengue virus DENV) genomes, we attempted to reconstruct the genome using tiled primers. For WNV we followed the procedure described in Quick et al. (2017)^1^ exactly, except that we used our previously isolated RNA (which we isolated with TriZol/chloroform rather than with a kit). For dengue virus, we followed the protocol described in Vogels et al. (2024)^2^. Additionally, we reconstructed the genomes of WNV isolated from two wildlife samples processed in our lab space that we suspected could be the source of the WNV sequences detected in our samples. We used primer pairs from <https://docs.google.com/spreadsheets/d/1zMfUv1IV5-Sy-AeOPhKHdoZMUxhD1R_RsB5gPswqx40/edit?gid=0#gid=0> for WNV, and the primer pairs for DENV-2 from <https://www.protocols.io/view/dengueseq-a-pan-serotype-whole-genome-amplicon-seq-kqdg39xxeg25/v3/materials> (38 pairs for West Nile virus, 37 pairs for DENV-2) designed to capture the entirety of each genome in overlapping ~ 400 bp sections. Primers were multiplexed into two pools, such that primers amplifying adjacent portions of the genome were kept in separate pools. Following amplicon generation and cleanup, we prepared libraries with the native barcoding protocol and sequenced on the Oxford Nanopore Technologies PromethION sequencer (WNV) and submitted pooled cDNA amplicon libraries to PlasmidSaurus for nanopore sequencing (DENV). We generated consensus sequences using the *artic* ‘field-bioinformatics’(v. 1.6.2) pipeline (<https://github.com/artic-network/fieldbioinformatics>) with a 20 X depth threshold to perform read filtering, primer trimming, and amplicon coverage normalization. We used megaBLAST^3^ to search for genomes with high levels of similarity to our consensus genomes.

**Supplemental Results**

*Batch correction of gottcha2-generated data*

RNA extraction batch 17 (B17) was determined to be the most effective reference batch for performing batch correction with the default method in package *ConQuR*^4^ , and allowed us to reduce the effect of RNA extraction batch on the variation in our data by 9.9% (Table S2, Figure S1).

**Table S1.** We tested the default and penalized algorithms for batch correction using each RNA extraction batch as a reference against which other batches were compared. Using the penalized algorithm with RNA extraction batch B17 as a reference was the most effective (bold).

| **RNA extraction batch ID** | **Method** | **Batch R^2^** | **Batch Reduction (%)** |
| --- | --- | --- | --- |
| Original | None | 0.100 | 0.0 |
| B01 | Default | 0.114 | -13.6 |
| B01 | Penalized | 0.091 | 9.1 |
| B02 | Default | 0.081 | 19.6 |
| B02 | Penalized | 0.085 | 14.9 |
| B03 | Default | 0.125 | -24.5 |
| B03 | Penalized | 0.089 | 11.0 |
| B04 | Default | 0.151 | -50.0 |
| B04 | Penalized | 0.086 | 14.5 |
| B05 | Default | 0.133 | -32.5 |
| B05 | Penalized | 0.089 | 11.0 |
| B06 | Default | 0.091 | 9.3 |
| B06 | Penalized | 0.085 | 15.2 |
| B07 | Default | 0.359 | -258.1 |
| B07 | Penalized | 0.127 | -26.5 |
| B08 | Default | 0.115 | -14.8 |
| B08 | Penalized | 0.095 | 5.1 |
| B09 | Default | 0.086 | 14.7 |
| B09 | Penalized | 0.096 | 4.4 |
| B10 | Default | 0.108 | -7.5 |
| B10 | Penalized | 0.105 | -4.6 |
| B11 | Default | 0.337 | -235.7 |
| B11 | Penalized | 0.095 | 4.9 |
| B12 | Default | 0.088 | 12.2 |
| B12 | Penalized | 0.090 | 10.7 |
| B13 | Default | 0.178 | -77.4 |
| B13 | Penalized | 0.085 | 15.5 |
| B14 | Default | 0.178 | -77.3 |
| B14 | Penalized | 0.085 | 14.9 |
| B15 | Default | 0.194 | -93.5 |
| B15 | Penalized | 0.089 | 11.5 |
| B16 | Default | 0.104 | -3.3 |
| B16 | Penalized | 0.082 | 18.3 |
| B17 | Default | 0.105 | -4.2 |
| **B17** | **Penalized** | **0.080** | **19.8** |
| B18 | Default | 0.119 | -18.4 |
| B18 | Penalized | 0.105 | -4.7 |
| B19 | Default | 0.089 | 11.8 |
| B19 | Penalized | 0.149 | -48.0 |

**Table S2.** Bacterial reads from potentially pathogenic bacteria present in samples at read counts greater than or equal to 10 and contig counts greater than or equal to two. All reference genomes to which contigs had highest-similarity alignments are listed. Entries are listed in descending order of positions covered and total reads per species in a sample.

| **Sample** | **Species** | **TAXID** | **Reference(s)** | **Total Reads** | **N Contigs** | **Positions Covered** | **Mean Depth** | **Max Depth** |
| --- | --- | --- | --- | --- | --- | --- | --- | --- |
| DK19 | *Acinetobacter guillouiae* | 106649 | NZ_AP014630.1, NZ_CP058272.1, NZ_CP083989.1, NZ_CP094620.1, NZ_CP118892.1 | 20 | 5 | 5,005 | 0.00 | 4 |
| DK20 | *Morganella morganii* | 582 | NZ_CP014026.2, NZ_CP026651.1, NZ_CP027177.1, NZ_CP061513.1, NZ_CP070407.1, NZ_CP128435.1, NZ_CP132323.1, NZ_LR699007.1 | 20 | 8 | 3,110 | 0.00 | 3 |
| DK29 | *Acinetobacter haemolyticus* | 29430 | NZ_CP018260.1, NZ_CP116040.1 | 20 | 2 | 1,564 | 0.00 | 10 |
| DK38 | *Enterobacter kobei* | 208224 | NZ_CP083829.1, NZ_CP088229.1 | 20 | 2 | 1,402 | 0.00 | 20 |
| F19 | *Enterobacter kobei* | 208224 | NZ_CP017181.1, NZ_CP083862.1, NZ_CP088229.1, NZ_CP096849.1 | 20 | 4 | 4,124 | 0.00 | 12 |
| F34 | *Klebsiella variicola* | 244366 | NZ_CP060807.1, NZ_CP077805.1 | 20 | 2 | 2,542 | 0.00 | 9 |
| F35 | *Enterobacter kobei* | 208224 | NZ_CP083862.1, NZ_CP088229.1, NZ_CP096849.1 | 20 | 3 | 2,975 | 0.00 | 15 |
| F45 | *Acinetobacter guillouiae* | 106649 | NZ_AP014630.1, NZ_CP058272.1, NZ_CP083989.1, NZ_CP094620.1, NZ_CP118892.1 | 20 | 5 | 5,229 | 0.00 | 5 |
| F47 | *Citrobacter sedlakii* | 67826 | NZ_CP057611.1, NZ_CP057620.1 | 20 | 2 | 1,302 | 0.00 | 13 |
| F52 | *Aeromonas hydrophila* | 644 | NZ_CP064382.1, NZ_LR963141.1 | 20 | 2 | 1,631 | 0.00 | 18 |
| F52 | *Vibrio cholerae* | 666 | NZ_LT897797.1, NZ_LT992490.1 | 20 | 2 | 4,686 | 0.00 | 7 |
| F75 | *Streptococcus salivarius* | 1304 | NZ_CP015282.1, NZ_CP015283.1, NZ_CP053998.1, NZ_CP054153.1, NZ_CP090007.1, NZ_CP133476.1 | 20 | 6 | 1,605 | 0.00 | 7 |
| F75 | *Streptococcus intermedius* | 1338 | NZ_AP014880.1, NZ_CP020433.2, NZ_CP053999.1, NZ_LS483436.1 | 20 | 4 | 1,415 | 0.00 | 10 |
| F75 | *Gemella haemolysans* | 1379 | NZ_CP050965.1, NZ_LR134484.1 | 20 | 2 | 3,242 | 0.00 | 5 |
| F75 | *Rothia aeria* | 172042 | NZ_CP068102.1, NZ_LR134479.1 | 20 | 2 | 2,574 | 0.00 | 6 |
| F75 | *Staphylococcus equorum* | 246432 | NZ_CP012968.1, NZ_CP013114.1, NZ_CP013714.1, NZ_CP013980.1, NZ_CP093841.1 | 20 | 5 | 3,476 | 0.00 | 11 |
| F75 | *Prevotella melaninogenica* | 28132 | NZ_CP022040.2, NZ_CP022041.2, NZ_CP085940.1, NZ_CP085941.1, NZ_CP085942.1, NZ_CP085943.1 | 20 | 6 | 1,837 | 0.00 | 6 |
| F75 | *Veillonella rogosae* | 423477 | NZ_CP110418.1, NZ_CP117967.1 | 20 | 2 | 1,170 | 0.00 | 9 |
| FAL23 | *Enterobacter kobei* | 208224 | NZ_CP083829.1, NZ_CP083862.1, NZ_CP088229.1, NZ_CP096849.1 | 20 | 4 | 2,430 | 0.00 | 15 |
| FAL52 | *Enterococcus faecalis* | 1351 | NZ_AP018538.1, NZ_AP018543.1, NZ_CP042213.2, NZ_CP045598.1, NZ_CP046022.1, NZ_CP065317.1, NZ_CP071176.1, NZ_CP082231.1, NZ_CP092907.1, NZ_CP103863.1, NZ_CP116571.1, NZ_CP120228.1, NZ_LR698844.1, NZ_OD940420.1, NZ_OD940422.1 | 20 | 15 | 4,455 | 0.00 | 7 |
| FMS115 | *Citrobacter youngae* | 133448 | NZ_CP049738.1, NZ_CP049739.1, NZ_CP102500.1 | 20 | 3 | 5,929 | 0.00 | 5 |
| FMS116 | *Enterobacter kobei* | 208224 | NZ_CP088229.1, NZ_CP104726.1 | 20 | 2 | 14,665 | 0.00 | 2 |
| FMS116 | *Escherichia albertii* | 208962 | NZ_CP043271.1, NZ_CP117637.1 | 20 | 2 | 1,987 | 0.00 | 23 |
| FMS116 | *Pantoea agglomerans* | 549 | NZ_CP059089.1, NZ_OW970315.1 | 20 | 2 | 6,337 | 0.00 | 6 |
| FMS116 | *Pantoea ananatis* | 553 | NZ_CP059084.1, NZ_CP059085.1, NZ_CP090356.1, NZ_CP099535.1 | 20 | 4 | 5,894 | 0.00 | 4 |
| FMS116 | *Citrobacter sedlakii* | 67826 | NZ_CP057611.1, NZ_CP057620.1, NZ_CP071070.1 | 20 | 3 | 3,585 | 0.00 | 10 |
| FMS121 | *Aeromonas hydrophila* | 644 | NZ_AP023398.1, NZ_CP124746.1 | 20 | 2 | 2,047 | 0.00 | 18 |
| FMS132 | *Klebsiella variicola* | 244366 | NZ_CP060807.1, NZ_CP077805.1 | 20 | 2 | 3,228 | 0.00 | 13 |
| FMS136 | *Enterobacter kobei* | 208224 | NC_018405.1, NZ_AP022126.1, NZ_AP022431.1, NZ_AP022446.1, NZ_AP022498.1, NZ_CP015227.1, NZ_CP017181.1, NZ_CP032897.1, NZ_CP083828.1, NZ_CP088119.1, NZ_CP091481.1, NZ_CP110873.1 | 20 | 12 | 2,253 | 0.00 | 12 |
| FMS142 | *Acinetobacter guillouiae* | 106649 | NZ_AP014630.1, NZ_CP058272.1 | 20 | 2 | 3,397 | 0.00 | 15 |
| FMS145 | *Klebsiella variicola* | 244366 | NZ_CP060807.1, NZ_CP077805.1 | 20 | 2 | 3,638 | 0.00 | 9 |
| FMS149 | *Acinetobacter haemolyticus* | 29430 | NZ_CP018260.1, NZ_CP018871.1, NZ_CP031979.1, NZ_CP032002.1, NZ_CP038009.1, NZ_CP116040.1 | 20 | 6 | 2,016 | 0.00 | 15 |
| FMS189 | *Citrobacter youngae* | 133448 | NZ_CP014030.2, NZ_CP049739.1, NZ_CP102500.1, NZ_LR134485.1 | 20 | 4 | 886 | 0.00 | 20 |
| FMS192 | *Kosakonia cowanii* | 208223 | NZ_CP034225.1, NZ_CP069322.1 | 20 | 2 | 2,174 | 0.00 | 14 |
| FMS196 | *Morganella morganii* | 582 | NZ_CP043955.1, NZ_CP132323.1, NZ_LR699007.1 | 20 | 3 | 450 | 0.00 | 20 |
| FMS197 | *Enterobacter kobei* | 208224 | NZ_CP083862.1, NZ_CP096849.1, NZ_CP110873.1 | 20 | 3 | 7,735 | 0.00 | 6 |
| FMS197 | *Citrobacter sedlakii* | 67826 | NZ_CP057611.1, NZ_CP057620.1 | 20 | 2 | 2,847 | 0.00 | 12 |
| FMS197 | *Enterobacter cancerogenus* | 69218 | NZ_CP025225.1, NZ_LR881936.1 | 20 | 2 | 5,798 | 0.00 | 5 |
| FMS198 | *Enterobacter kobei* | 208224 | NZ_CP042578.1, NZ_CP043511.1, NZ_CP050073.1, NZ_CP083828.1, NZ_CP083857.1, NZ_CP083862.1, NZ_CP088229.1, NZ_CP096849.1, NZ_CP104724.1 | 20 | 9 | 11,845 | 0.00 | 2 |
| FMS198 | *Acinetobacter haemolyticus* | 29430 | NZ_CP018871.1, NZ_CP018873.1, NZ_CP030880.1, NZ_CP031979.1, NZ_CP116040.1 | 20 | 5 | 7,543 | 0.00 | 13 |
| FMS87 | *Enterobacter kobei* | 208224 | NZ_CP083828.1, NZ_CP083829.1 | 20 | 2 | 7,096 | 0.00 | 5 |
| FMS87 | *Citrobacter amalonaticus* | 35703 | NZ_CP057631.1, NZ_CP057633.1 | 20 | 2 | 1,606 | 0.05 | 13 |
| FMS93 | *Pseudomonas mosselii* | 78327 | NZ_CP023299.1, NZ_CP128544.1 | 20 | 2 | 3,062 | 0.00 | 10 |
| FMS94 | *Enterobacter kobei* | 208224 | NZ_CP083862.1, NZ_CP088229.1, NZ_CP096849.1 | 20 | 3 | 2,975 | 0.00 | 13 |
| FMS94 | *Citrobacter sedlakii* | 67826 | NZ_CP057611.1, NZ_CP057620.1 | 20 | 2 | 2,260 | 0.00 | 11 |
| FMS95 | *Enterobacter kobei* | 208224 | NZ_AP022126.1, NZ_CP017181.1, NZ_CP075345.1, NZ_CP083862.1, NZ_CP096849.1 | 20 | 5 | 2,263 | 0.00 | 16 |
| FMS99 | *Vibrio furnissii* | 29494 | NC_016602.1, NZ_CP040990.1, NZ_CP046797.1, NZ_CP051103.1, NZ_CP064379.1, NZ_CP089603.1, NZ_CP100422.1, NZ_CP119521.1, NZ_CP128200.1 | 20 | 9 | 1,678 | 0.00 | 20 |
| F52 | *Shewanella algae* | 38313 | NZ_CP047422.1, NZ_CP055159.1, NZ_CP068230.1 | 19 | 3 | 2,425 | 0.00 | 13 |
| F75 | *Streptococcus oralis* | 1303 | NC_015291.1, NZ_AP018338.1, NZ_CP016207.1, NZ_CP029257.1, NZ_CP054134.1, NZ_CP066172.1, NZ_CP069427.1, NZ_CP079724.1 | 19 | 8 | 1,617 | 0.00 | 9 |
| F75 | *Streptococcus mitis* | 28037 | NC_013853.1, NZ_CP012646.1, NZ_CP028414.1, NZ_CP028415.1, NZ_CP067992.1, NZ_CP077259.1, NZ_CP133471.1 | 19 | 7 | 1,556 | 0.00 | 8 |
| DK29 | *Bordetella trematum* | 123899 | NZ_CP016340.1, NZ_CP018898.1, NZ_CP036357.1, NZ_CP049957.1, NZ_LT546645.1 | 18 | 5 | 2,694 | 0.00 | 5 |
| F53 | *Achromobacter spanius* | 217203 | NZ_CP034689.1, NZ_CP079940.1 | 18 | 2 | 5,091 | 0.00 | 3 |
| F75 | *Staphylococcus caprae* | 29380 | NZ_AP018585.1, NZ_AP018587.1, NZ_CP031271.1, NZ_CP051643.1 | 18 | 4 | 741 | 0.00 | 8 |
| F75 | *Pantoea eucrina* | 472693 | NZ_CP083448.1, NZ_CP083449.1, NZ_CP083450.1 | 18 | 3 | 1,550 | 0.00 | 8 |
| FMS146 | *Achromobacter spanius* | 217203 | NZ_CP025030.1, NZ_CP032084.1, NZ_CP034689.1, NZ_CP079940.1, NZ_LR134302.1 | 18 | 5 | 3,071 | 0.00 | 4 |
| F27 | *Pseudomonas mosselii* | 78327 | NZ_CP104107.1, NZ_CP128544.1 | 17 | 2 | 1,542 | 0.00 | 9 |
| DK20 | *Weissella confusa* | 1583 | NZ_CP027563.1, NZ_CP027565.1, NZ_CP049097.1, NZ_CP110106.1, NZ_CP120516.1 | 16 | 5 | 2,667 | 0.00 | 3 |
| DK63 | *Morganella morganii* | 582 | NC_020418.1, NZ_CP014026.2, NZ_CP025933.1, NZ_CP026046.1, NZ_CP032295.1, NZ_CP034944.1, NZ_CP043955.1, NZ_CP064830.1, NZ_CP068145.1, NZ_LR699007.1 | 16 | 10 | 2,749 | 0.00 | 8 |
| F75 | *Enterococcus gilvus* | 160453 | NZ_CP030932.1, NZ_CP030933.1 | 16 | 2 | 2,621 | 0.00 | 6 |
| F75 | *Cutibacterium modestum* | 2559073 | NZ_AP024747.1, NZ_AP024748.1, NZ_CP017040.1, NZ_CP017041.1 | 16 | 4 | 2,500 | 0.00 | 6 |
| FMS116 | *Pantoea eucrina* | 472693 | NZ_CP083448.1, NZ_CP083450.1 | 16 | 2 | 5,321 | 0.00 | 3 |
| DK19 | *Gemella haemolysans* | 1379 | NZ_CP050965.1, NZ_CP083637.1 | 15 | 2 | 5,813 | 0.00 | 4 |
| F24 | *Burkholderia mallei* | 13373 | NZ_CP009535.1, NZ_CP009551.1, NZ_CP018373.1, NZ_CP019042.1, NZ_CP025302.1, NZ_CP071757.1, NZ_CP116601.1, NZ_LR595894.1, NZ_LR595896.1 | 15 | 9 | 5,967 | 0.00 | 2 |
| F75 | *Lactobacillus gasseri* | 1596 | NC_008530.1, NZ_CP006803.1, NZ_CP049762.1, NZ_CP072178.1, NZ_CP087761.1, NZ_CP090409.1 | 15 | 6 | 1,432 | 0.00 | 5 |
| F75 | *Haemophilus haemolyticus* | 726 | NZ_CP027235.1, NZ_CP031243.1 | 15 | 2 | 1,140 | 0.00 | 8 |
| FMS94 | *Bacillus thuringiensis* | 1428 | NC_018486.1, NC_018500.1, NZ_CP010577.1, NZ_CP014287.1, NZ_CP014847.1, NZ_CP032613.1, NZ_CP037890.1, NZ_CP045030.1, NZ_CP070339.1, NZ_CP071743.1, NZ_CP076539.1, NZ_CP095769.1, NZ_CP114399.1, NZ_CP133375.1 | 15 | 14 | 3,351 | 0.00 | 2 |
| FAL92 | *Lactococcus garvieae* | 1363 | NC_017490.1, NZ_AP026069.1, NZ_CP026502.1, NZ_CP071285.1, NZ_CP084377.1, NZ_CP118627.1, NZ_CP118950.1 | 14 | 7 | 860 | 0.00 | 14 |
| FMS147 | *Burkholderia mallei* | 13373 | NZ_AP028073.1, NZ_AP028077.1, NZ_CM003194.1, NZ_CP018373.1, NZ_CP019042.1, NZ_CP025306.1, NZ_CP041218.1, NZ_CP041220.1, NZ_CP073731.1, NZ_CP073734.1, NZ_JAGTPE010000001.1, NZ_LR595896.1 | 14 | 12 | 8,616 | 0.00 | 3 |
| FMS95 | *Citrobacter sedlakii* | 67826 | NZ_CP057611.1, NZ_CP057620.1, NZ_CP071070.1 | 14 | 3 | 2,804 | 0.00 | 5 |
| F19 | *Achromobacter spanius* | 217203 | NZ_CP034689.1, NZ_CP079940.1 | 13 | 2 | 3,341 | 0.00 | 4 |
| F4 | *Acinetobacter haemolyticus* | 29430 | NZ_CP018873.1, NZ_CP030880.1, NZ_CP031979.1, NZ_CP031984.1, NZ_CP116040.1 | 13 | 5 | 8,091 | 0.00 | 3 |
| F75 | *Haemophilus parainfluenzae* | 729 | NZ_CP031477.1, NZ_CP063122.1, NZ_CP065991.1, NZ_CP125087.1, NZ_CP133470.1 | 13 | 5 | 2,234 | 0.00 | 5 |
| FMS124 | *Klebsiella variicola* | 244366 | NZ_CP060807.1, NZ_CP077805.1 | 13 | 2 | 2,790 | 0.00 | 6 |
| F39 | *Achromobacter spanius* | 217203 | NZ_CP034689.1, NZ_CP079940.1 | 12 | 2 | 3,496 | 0.00 | 5 |
| F75 | *Corynebacterium coyleae* | 53374 | NZ_CP047198.1, NZ_CP077302.1, NZ_CP083648.1 | 12 | 3 | 862 | 0.00 | 3 |
| F75 | *Haemophilus parahaemolyticus* | 735 | NZ_CP038817.1, NZ_CP069510.1 | 12 | 2 | 620 | 0.00 | 9 |
| FAL86 | *Prevotella nigrescens* | 28133 | NZ_CP072342.1, NZ_CP082843.1, NZ_CP133464.1 | 12 | 3 | 359 | 0.00 | 12 |
| FMS106 | *Pseudomonas luteola* | 47886 | NZ_CP033105.1, NZ_CP044086.1 | 12 | 2 | 198 | 0.00 | 12 |
| FMS144 | *Citrobacter sedlakii* | 67826 | NZ_CP057611.1, NZ_CP057620.1, NZ_CP071070.1 | 12 | 3 | 1,267 | 0.00 | 5 |
| FMS189 | *Enterobacter kobei* | 208224 | NZ_AP022431.1, NZ_CP032897.1, NZ_CP048736.1, NZ_CP071178.1, NZ_CP088119.1, NZ_CP097572.1 | 12 | 6 | 735 | 0.00 | 5 |
| DK19 | *Streptococcus pneumoniae* | 1313 | NC_003028.3, NZ_AP018936.1, NZ_CM001835.1, NZ_CP025076.1, NZ_CP025256.1, NZ_CP038252.1, NZ_CP071871.1, NZ_CP091450.1, NZ_LN831051.1 | 11 | 9 | 1,312 | 0.00 | 5 |
| DK42 | *Acinetobacter haemolyticus* | 29430 | NZ_CP018260.1, NZ_CP018871.1, NZ_CP031984.1, NZ_CP031991.1, NZ_CP038009.1, NZ_CP116040.1 | 11 | 6 | 2,022 | 0.00 | 4 |
| F75 | *Actinomyces oris* | 544580 | NZ_AP025590.1, NZ_CP014232.1, NZ_CP040005.1, NZ_CP066060.1, NZ_CP068012.1, NZ_CP116224.1 | 11 | 6 | 537 | 0.00 | 6 |
| FMS121 | *Aeromonas dhakensis* | 196024 | NZ_CP077394.1, NZ_CP084349.1, NZ_CP102368.1, NZ_CP121801.1, NZ_CP121802.1, NZ_LR963089.1, NZ_LR963112.1, NZ_LR963122.1 | 11 | 8 | 2,607 | 0.00 | 4 |
| FMS198 | *Citrobacter koseri* | 545 | NZ_AP023452.1, NZ_OW969691.1 | 11 | 2 | 1,342 | 0.00 | 7 |
| DK44 | *Acinetobacter haemolyticus* | 29430 | NZ_CP018260.1, NZ_CP018871.1, NZ_CP031972.1, NZ_CP031991.1, NZ_CP038009.1, NZ_CP116040.1 | 10 | 6 | 1,109 | 0.00 | 5 |
| F21 | *Pseudomonas luteola* | 47886 | NZ_CP033105.1, NZ_CP044086.1 | 10 | 2 | 618 | 0.00 | 6 |
| F35 | *Citrobacter sedlakii* | 67826 | NZ_CP057611.1, NZ_CP057620.1, NZ_CP071070.1 | 10 | 3 | 2,030 | 0.00 | 6 |
| F60 | *Acinetobacter guillouiae* | 106649 | NZ_AP014630.1, NZ_CP058272.1, NZ_CP083989.1, NZ_CP094620.1 | 10 | 4 | 1,813 | 0.00 | 9 |
| F75 | *Actinomyces naeslundii* | 1655 | NZ_CP066049.1, NZ_CP113787.1 | 10 | 2 | 832 | 0.00 | 6 |
| F75 | *Anaerococcus vaginalis* | 33037 | NZ_CP066014.1, NZ_CP067016.1 | 10 | 2 | 1,949 | 0.00 | 3 |
| FA17 | *Weissella confusa* | 1583 | NZ_CP049097.1, NZ_CP080582.1, NZ_CP110106.1 | 10 | 3 | 1,792 | 0.00 | 5 |
| FAL3 | *Listeria monocytogenes* | 1639 | NC_018587.1, NZ_CP011345.1, NZ_CP019622.1, NZ_CP050029.1, NZ_CP058256.1, NZ_CP087264.1 | 10 | 6 | 640 | 0.00 | 5 |
| FAL72 | *Corynebacterium propinquum* | 43769 | NZ_CP068160.1, NZ_CP068161.1, NZ_CP100361.1, NZ_CP100363.1, NZ_CP100364.1, NZ_CP100371.1 | 10 | 6 | 3,076 | 0.00 | 3 |
| FAL9 | *Vibrio cholerae* | 666 | NZ_LT897797.1, NZ_LT992490.1 | 10 | 2 | 2,842 | 0.00 | 3 |
| FMS142 | *Klebsiella variicola* | 244366 | NZ_CP060807.1, NZ_CP077805.1 | 10 | 2 | 3,128 | 0.00 | 6 |

**Table S3.** Summary of RDT and metagenomic sequencing results. Percentages calculated from total samples with corresponding sequencing data analyzed by CZ ID (n = 359). Sequencing detection based on CZID pathogen identification with Z-score filtering.

| **RDT Results** | | | **Sequencing Results** | | **Concordance** | | **Discordance** | | **RDT Unknown** | |
| --- | --- | --- | --- | --- | --- | --- | --- | --- | --- | --- |
| **RDT+ n (%)** | **RDT- n (%)** | **RDT Unknown n (%)** | **Seq+ n (%)** | **Seq- n (%)** | **Both+ n** | **Both- n** | **RDT+/Seq- n** | **RDT-/Seq+ n** | **RDT?/Seq+ n** | **RDT?/Seq- n** |
| 21 (5.8) | 118 (32.9) | 220 (61.3) | 11 (3.1) | 348 (96.9) | 1 | 114 | 20 | 4 | 6 | 214 |

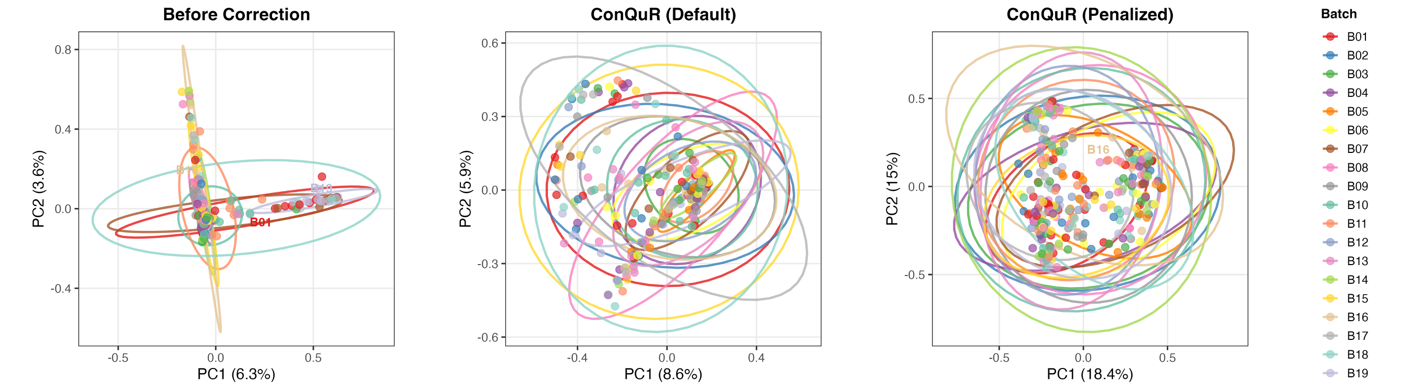

**Figure S1.** PCoAs based on Bray-Curtis dissimilarities of samples colored based on RNA extraction batch ID (A) prior to batch correction, (B) after default batch correction using ConQuR and Batch 17 as a reference, and (C) after penalized batch correction using ConQuR and Batch 17 as a reference.

*PrimalSeq and DengueSeq*

We generated consensus WNV genomes from 11 patient samples and four wildlife-derived isolates, and one blank (which became contaminated during lab work). We used mafft with default (auto) settings^5^ to perform a multiple sequence alignment with all consensus WNV sequences we generated, 2,267 North American WNV sequences pulled from the North American WNV project on Next Strain (<https://nextstrain.org/WNV/NA>), and 49 NCBI-accessioned sequences identified by a review of WNV genomic surveillance gaps in Africa ^6^. After multiple sequence alignment, we built a phylogenetic tree using iqtree3 ^7^ with 1000 bootstraps. All patient-derived sequences clustered together within a clade of other North American sequences and were determined to be contamination from the wildlife-derived isolates (Figure S2). Wildlife-derived WNV sequences are archived in GenBank with accession IDs PX462748, and PX462749. We were only able to construct partial consensus DENV-2 genomes from two of the three putative DENV-positive samples (most amplicons were not generated and in one sample, no amplicons were successfully generated). The successfully amplified regions of the DENV-2 genome were also determined to be lab contamination; they had 100 % identity to a laboratory strain of DENV-2 (GenBank accession JN796245.1).

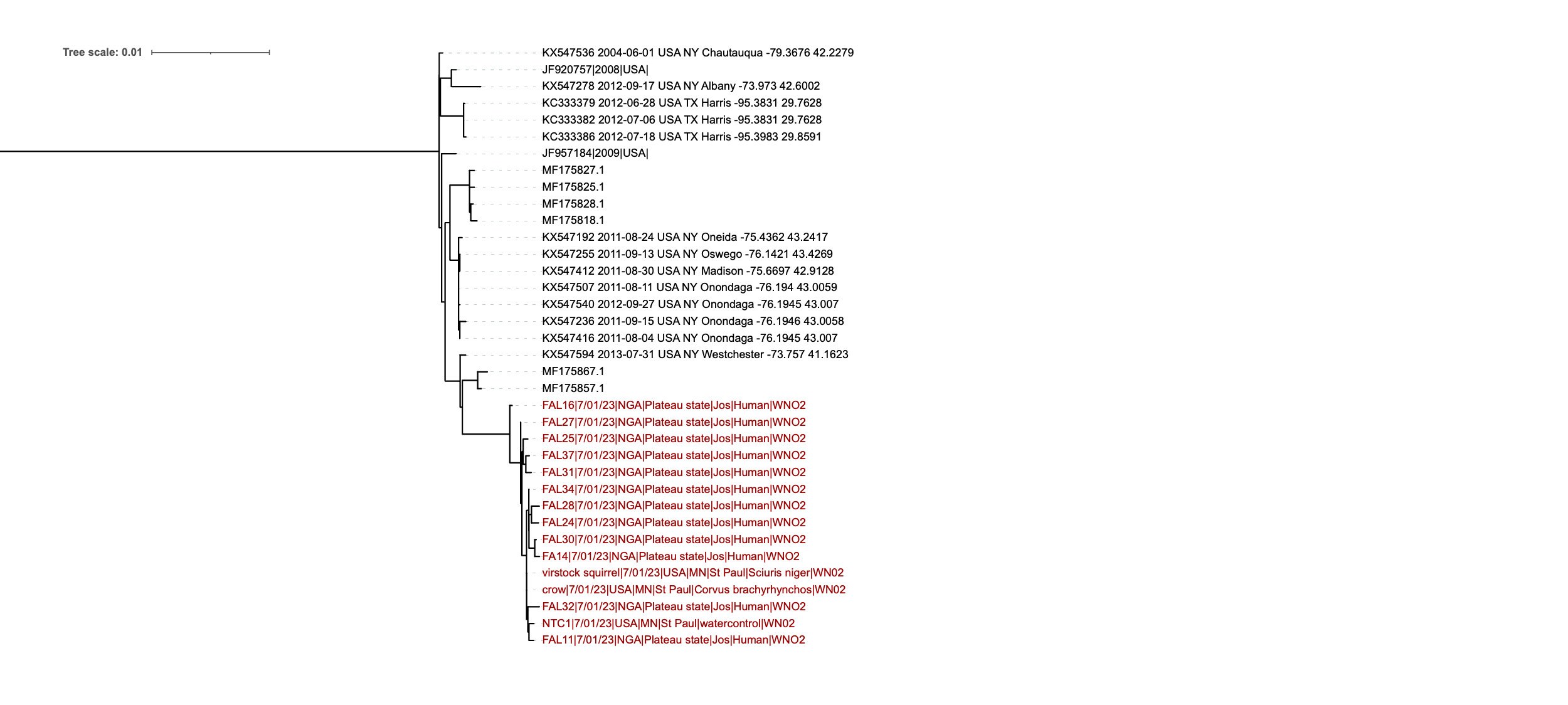

**Figure S2.** Pruned maximum-likelihood tree of WNV sequences generated in our study (red) and the most closely related sequences from North America (most leaves omitted for readability).

7.	Wong, Thomas K.F., Ly-Trong, Nhan, Ren, Huaiyan. IQ-TREE 3: Phylogenomic Inference Software using Complex Evolutionary Models
